# Holistic meta-analysis of *Caenorhabditis elegans* germ granule proteomics reveals complex dynamics and new candidate granule associated proteins

**DOI:** 10.64898/2026.03.18.712568

**Authors:** Carlotta Wills, Alyson Ashe

## Abstract

Spatiotemporal organisation of biological molecules is a key driver of cellular processes, including many post-transcriptional epigenetic processes. The germline-specific germ granules are biomolecular condensates that act as hubs for mRNA and small RNA processing and are core regulators of germline gene expression programming. Germ granules have been studied extensively in *C. elegans*, and recent developments have led to many subdivisions of the germ granule into specialised compartments. Rapid advancements in microscopy and protein-protein interaction (PPI) screening techniques have produced a large amount of data towards characterising the localisation of proteins to specific granules. However, common methods used to probe PPIs are limited in their ability to robustly detect valid interactions, especially the multivalent and sometimes transient ones observed in granule environments.

Here we perform a meta-analysis of granule protein interaction screens. While these experiments generally enrich for proteins matching the profile of granule-associated proteins, we find that when considering screens individually, reproducibility is surprisingly low, highlighting not only the variability inherent in these methods but also the dynamic nature of the PPI networks present in granules. We developed an algorithm to provide a measure of each proteins’ association with specific granules across various experiments. By further clustering and investigation of the resulting score matrix, we demonstrate the power of this holistic approach to provide deeper insights into germ granule organisation and highlight novel can provide a resource to better inform future investigations into granules and their constituent proteins.

## Introduction

The coordination of biochemical processes within cells depends on spatiotemporal organisation of the necessary molecular components. Many such processes occur within membraneless condensates (or “granules”), which coalesce by liquid-liquid phase separation (LLPS) (Banani et al. 2017). The liquid-like nature of these granules allows for them to dynamically and rapidly reorganise themselves in response to environmental or developmental signals. Formation and organisation of granules is largely dependent on multivalent interactions between the biomolecules residing within them: granule-associated proteins often contain LLPS-promoting intrinsically disordered regions (Nott et al. 2015), and RNA (Zhang et al. 2015).

The germline has specialised ribonucleoprotein (RNP) granules (here, referred to collectively as “germ granules”). These granules may be referred to by many names, are implicated in germ cell development and specification (Thomas et al. 2023), and act as a hub for germline-critical small RNA pathways (Ouyang and Seydoux 2022). In mammals, germ granules in male germ cells are known as “intermitochondrial cement” or “chromatoid bodies”, depending on developmental stage, and have been shown to harbour piRNAs and their biogenesis machinery (Lehtiniemi and Kotaja 2018). Likewise, the “nuage” of germ cells in *Drosophila* are home to a number of Argonaute proteins amongst other RNA-binding proteins and are critical for production and amplification of piRNAs (Lin et al. 2023). The *Caenorhabditis elegans* germ granules are similarly home to several piRNA factors, and are important for other small RNA pathways such as siRNA biogenesis (Ouyang and Seydoux 2022). Despite the conservation of germ granules and their crucial involvement in germline processes, their precise mechanisms, organisation and architecture are poorly understood.

*C. elegans* is a valuable and prolific model in probing germ granule architecture and function. In *C. elegans*, the perinuclear germ granules are organised into various compartments. The first identified compartment was the P granule, named for its segregation into the P cell in the early embryo (Strome and Wood 1982). Its components include the Vasa homologs GLH-1/2 (Brangwynne et al. 2009) as well as the PGL class of proteins (Hanazawa et al. 2011). Situated over nuclear pores, P granules capture nascent mRNA as it exits the nucleus (Pitt et al. 2000; Schisa et al. 2001) and act as a hub for small RNA pathways (Ouyang et al. 2019; Dodson and Kennedy 2019). The P granules were long treated as synonymous with germ granules, but in recent years the characterisation of other compartments has advanced rapidly.

The subdivision of the germ granules began with the identification of the Mutator foci, where a number of siRNA biogenesis factors reside (Uebel et al. 2018). The Z granule was subsequently identified; its components include the zinc-finger protein ZNFX-1 (Ishidate et al. 2018; Wan et al. 2018), the Argonaute WAGO-4 (Wan et al. 2018) and the LOTUS and Tudor domain protein LOTR-1 (Marnik et al. 2022). The fourth identified compartment was the SIMR focus, which mediates the piRNA pathway and secondary siRNA biogenesis (Manage et al. 2020). Most recently, two more compartments of germ granules have been identified: the E granule required for the production of a specialised subset of siRNAs (Chen et al. 2024b) and the D granule positioned between nuclear pores and P granules (Lu et al. 2025).

Outside of the germ granules, there are other phase-separating granules of interest in the *C. elegans* germline. A sperm-specific granule known as the PEI granule has been identified as playing a role in the transmission of paternal epigenetic information (Schreier et al. 2022). Processing bodies (P bodies) are found in both somatic and germ cells, but play a distinct role in the germline where they interact with germ granules and assist in their compartmentalisation (Gallo et al. 2008; Du et al. 2023). Another granule found in both the soma and the germline is the stress granule, which as suggested by the name forms under various cellular stresses in the cytoplasm and is thought to have a protective function (Jud et al. 2008; Huelgas-Morales et al. 2016). A summary of granule organisation and known components can be found in Figure 1 and Supplemental Table S1. Notably, several of the currently validated granule-associated proteins interact in some way with RNA (e.g. as helicases, polymerases, or small RNA-binding Argonautes), highlighting the importance of these condensates in post-transcriptional processes.

**Figure 1.**
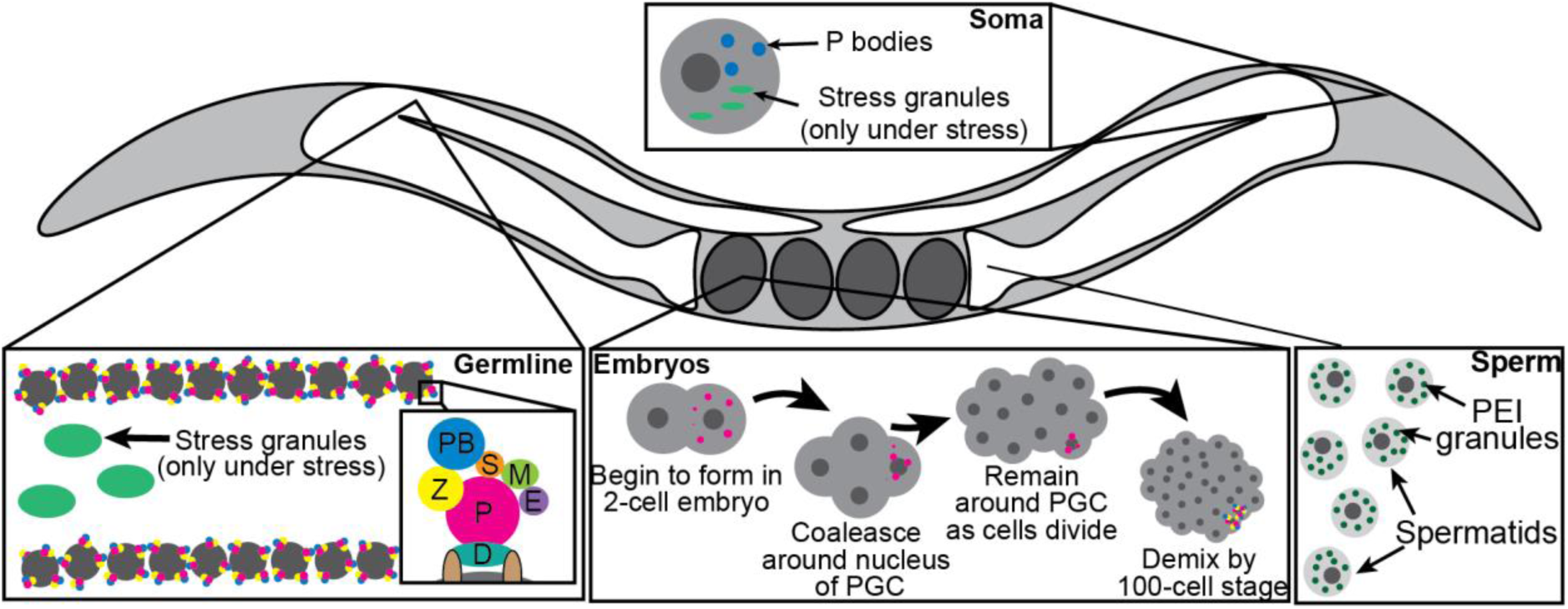
Dynamic RNP granules form in tissue-specific and temporal manners in *C. elegans*. In the germline (pachytene region represented here), nuclei are surrounded by germ granules, which sit over nuclear pores and subdivide into the P granule (P), Z granule (Z), Mutator foci (M), SIMR foci (S), E granule (E) and D granule (D). Germ granules can be observed in the primordial germ cells throughout embryogenesis. Also present in the germline are P bodies (PB), which sit adjacent to germ granules, and stress granules, which form in the cytoplasm only under conditions of cellular stress. P bodies and stress granules can also be observed in somatic cells. Sperm have a specialised granule known as the PEI granule.

An ongoing challenge in the study of granules is the definition of compartments and their constituents. Given their dynamic nature, defining proteins specific to a certain granule is highly challenging, especially as new developments lead to reassignment of previously discovered granule components. For example, the DEPS-1 protein was originally characterised as a constitutive P granule factor, and *deps-1* mutants display a P granule defective phenotype (Suen et al. 2020), however recent work using high-resolution imaging has redefined it to in fact be a Z granule component (Huang et al. 2024). Furthermore, it is apparent that granule components can move and shift depending on developmental stage or cellular condition. For example, P and Z granules are indistinguishable in the early embryo but segregate around the 100-cell stage (Wan et al. 2018), while the P body component CGH-1 (Jud et al. 2008) relocates to stress granules during cellular stress (Huelgas-Morales et al. 2016). Some granules themselves are also limited to particular cell-types, such as the PEI granule, which is only found in male sperm and harbours the Argonaute PPW-2 (also known as WAGO-3) (Schreier et al. 2022).

A common approach to studying granule location and formation is through high-resolution microscopy. Confocal microscopy is commonly employed to image granules through the use of known markers – for example, the imaging of PGL-1 either through endogenous fluorescent tagging (Wan et al. 2018) or immunofluorescence (Kawasaki et al. 1998) reveals the location and organisation of P granules. By tagging multiple proteins with distinct fluorescent tags, the organisation of different granules can be observed and the co-localisation of two proteins implicates them as belonging to the same compartment (Wan et al. 2018; Huang et al. 2024). However, the tagging of individual genes can be laborious, and requires a selected list of candidates to have already been identified. Furthermore, the ability to resolve different compartments varies highly depending on the microscopy technique used.

For larger-scale investigation of granule-associated proteins, researchers have often turned to protein-protein interaction (PPI) detection methods. These methods aim to identify interacting proteins of known granule components, with the implication being that proteins interacting with granule-associated proteins may localise to the same granule themselves. Early efforts to map PPIs in *C. elegans* employed the yeast two-hybrid (Y2H) method, first developed in the early 1990s to detect binary protein interactions *in vitro* (Chien et al. 1991; Walhout et al. 2002; Li et al. 2004). Such screens have provided useful data in mapping the protein “interactome”, but are still limited in their ability to detect valid *in vivo* interactions (Parrish et al. 2006).

In recent years, the use of mass spectrometry-coupled methods to detect interactions has allowed for more in-depth investigation of specific proteins’ interaction partners. The most common approach to identifying PPIs now involves affinity-purification of a target followed by mass spectrometry of the co-purifying proteins (AP-MS) (Dunham et al. 2012). Oftentimes, a tag that is used for imaging a protein by light microscopy – for example, GFP – can also be used for affinity purification, making AP-MS highly complementary with the microscopy methods often used to visualise granules. More recent developments of mass spectrometry-based PPI detection have involved the use of proximity labelling methods to better detect weak or transient interactions *in vivo* (Kim and Roux 2016).

Relative to other systems, there is limited publicly available proteomics data for *C. elegans*, but with growing adoption of these large-scale techniques, the pool of available data is expanding rapidly. As of February 2026, there are 50,342 proteomics datasets available on ProteomeXchange, a consortium of proteomics data repositories (Deutsch et al. 2023). Just 320 of these contain *C. elegans* data, with 84.4% of these uploaded in the current decade (2020-2026). Clearly, there is growing interest and demand for proteomics data in various *C. elegans* research fields.

While such data provide valuable insights into the interacting partners of their target proteins, individually they can only provide limited understanding of complex interaction networks like those found in granules. AP-MS, while targeted to a specific protein and robust in detecting strong interactions, ultimately removes a granule protein from its *in vivo* liquid-like context and may exclude weaker or transient interactions. Newer proximity-labelling techniques aim to address this limitation, but still only possess the ability to identify interactions for one protein at a time and may not capture the full spatiotemporal dynamics of granule interactomes.

Here, we review the current state of granule interactome research in *C. elegans*, focussing on the perinuclear granules that have been defined in the hermaphrodite germline (i.e. germ granule compartments and P bodies), as these granules have enough data to make a meta-analysis possible. We collate data from multiple publications that have performed large-scale experiments to identify potential interactors for known granule components. By comparing results across various experiments, we show that results from individual experiments are of variable reproducibility, suggesting that the complex and dynamic nature of granule proteomes demands more holistic approaches to their study. To this end, we perform a meta-analysis to synthesise available data across multiple granules and calculate association scores for several proteins across all granules examined. Through this, new insights into granule composition and organisation are gained, and future studies into “granule-omics” can be better informed.

## Results

### The growing base of granule interaction data

We identified 32 publications identifying candidate interactors using large-scale PPI screens (see Methods for full details), for a total of 51 unique datasets. Most of these used mass spectrometry to identify interaction partners for a protein of interest. Several of these studies were published within the past 5 years (Figure 2A). Of the studies that used mass spectrometry, 64% (18/28) also uploaded raw data to a proteomics repository, with the uploading of raw data becoming more common in recent years (Figure 2B).

**Figure 2.**
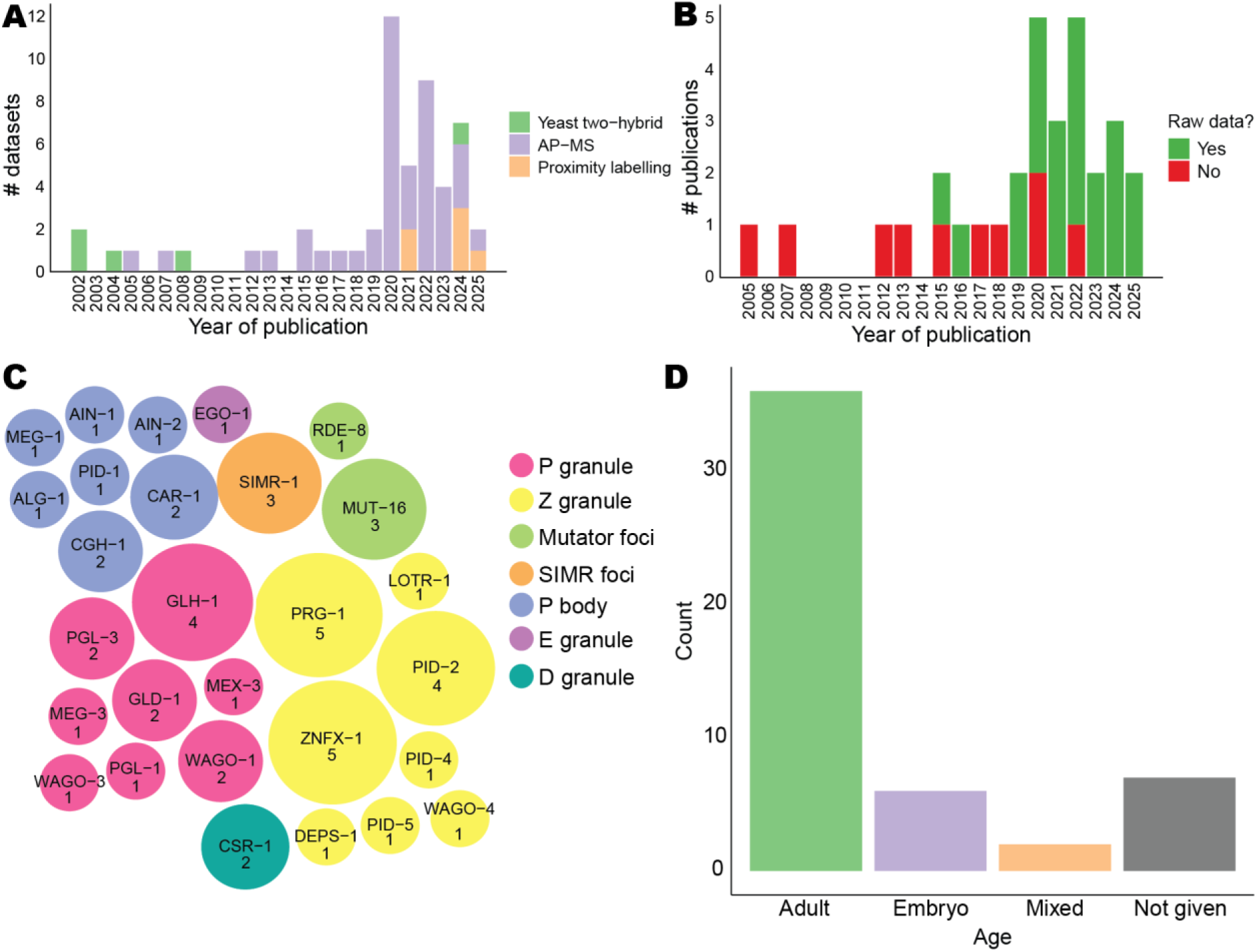
Overview of available datasets. **(A)** Summary of types of data in selected publications, organised by year and coloured by type of experiment. **(B)** Summary of availability of raw data for proteomics-based experiments. Raw data was scored as being available (“Yes”) if an accession number was provided in the publication, otherwise the data was scored as unavailable (“No”). **(C)** Number of datasets available from experiments targeted to specific proteins (untargeted Y2H screens not included), colour-coded by granule **(D)** Distribution of life stages used for experiments.

Aside from a handful of untargeted Y2H screens, most studies sought to identify interaction partners for specific proteins. The most studied granules in this context are the P granules, Z granules and P bodies (Figure 2C). Certain proteins have been subject to interaction screens more than others – such as the Piwi homolog PRG-1 and core Z granule component ZNFX-1, which have both been targeted 5 times. Unsurprisingly, the most recently identified granules (the E and D granules) are also the least studied. Most researchers chose to use synchronised adult populations for their experiments; however, some also chose to investigate embryos or use an unsynchronised mixed-stage population (Figure 2D).

### Germ granule interaction screens enrich for germline-related and RNA-binding proteins

In total, across all published screens examined, 5425 proteins were detected at least once, accounting for approximately 20% of the total reference proteome (5425/26696; total reference proteome count sourced from UniProt Proteome ID UP000001940). Many (>1500) of these proteins were only detected in a single experiment (Figure 3A). Proteins detected across several (> 20 experiments) were often known granule components, such as the P body protein PAB-1 (29 detections) and the P granule protein WAGO-1 (27 detections). Many ribosomal subunit proteins (RPL and RPS families) were also detected in high numbers.

**Figure 3.**
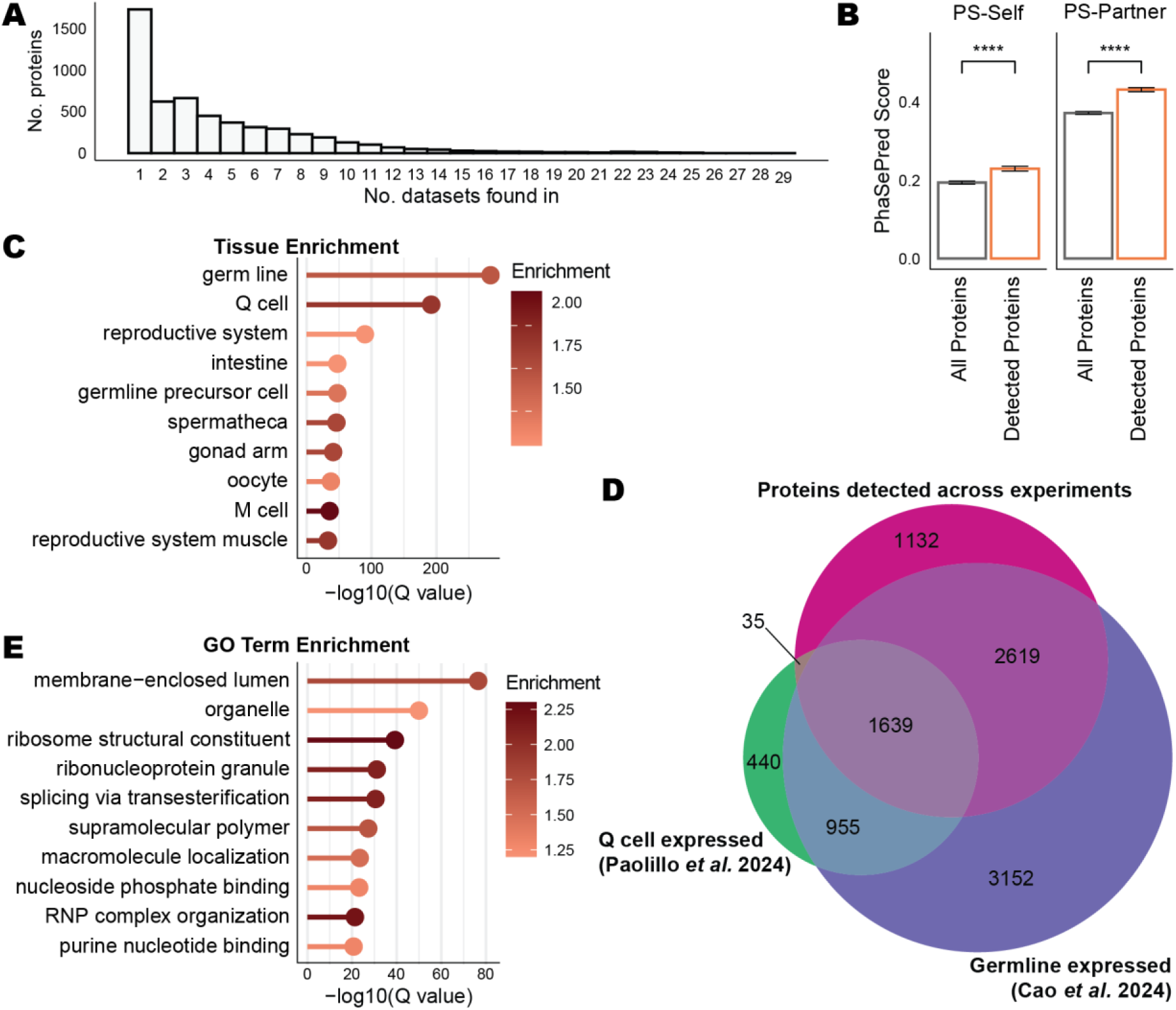
Proteins detected in granule-associated protein interaction screens fit the profile of other granule-associated proteins. **(A)** Histogram of number of detections for proteins identified using PPI screen methods. **(B)** Comparison of PhaSePred (Chen et al. 2022b) scores for self-assembly (PS-Self) and partner-dependent assembly (PS-Part) for all proteins in the PhaSePred database (n = 4143) and proteins detected across granule interaction screens present in the PhaSePred database (n = 1898). Bars show mean ± SEM, statistical comparison performed using *t* test. **** *p* < 0.0001. **(C)** Tissue enrichment analysis of all proteins detected across the datasets included in this study. **(D)** Overlap between all proteins detected across data included in this analysis, germline-expressed genes (Cao et al. 2024) and Q cell-expressed genes (Paolillo et al. 2024). **(E)** GO enrichment analysis of all proteins detected across the datasets included in this study. Both enrichment analyses were conducted using the WormBase Gene Set Enrichment Analysis tool (Angeles-Albores et al. 2016), and only the top twenty most significant enrichments (ranked by Q value) are shown.

Many efforts have been made to predict the likelihood of proteins to phase separate. The PhaSePred database (Chen et al. 2022b) collates predictive measures from various sources and provides an overall score of a protein’s propensity towards self-assembly (PS-Self) or partner-dependent assembly (PS-Part). Proteins detected in granule interaction screens show higher average PS-Self and PS-Part scores than the average of all *C. elegans* proteins in the PhaSePred database, indicating that these screens are likely enriching for proteins participating in LLPS as would be expected of granule-associated proteins (Figure 3B).

Given that many germ-granule localising proteins have heightened or even exclusive expression in the germline, we reason that interaction screens for these proteins should also enrich for other germline-expressed proteins. Indeed, enrichment analysis of all the proteins detected in these screens did reveal a strong enrichment for germline-associated proteins, as well as other specific germline-related tissues such as the oocyte, spermatheca, germline precursor cells and distal tip cells (Figure 3C). Curiously, the second-highest enrichment was for the Q cell, a neuroblast lineage unrelated to the germline. However, further examination revealed that most proteins detected as “Q cell” enriched sat within the large overlap between germline-expressed and Q cell-expressed genes, rather than being Q-cell specific proteins (Figure 3D).

GO enrichment analysis of the protein list further shows enrichment for terms relating to RNA binding and post-transcriptional processes, including a specific enrichment for “ribonucleoprotein granule” (Figure 3E). Alongside the tissue enrichment data, this suggests that these protein interaction screens are broadly successful at pulling down proteins of general relevance to germline and granule functions.

### Reproducibility of candidates from individual experiments is generally low

When reporting the results from a protein interaction screen, each experiment provides an independent list of candidate interactors. Candidates may be reported through either qualitative (e.g. the confidence score from a Y2H screen, or the presence/absence of a protein in a control vs targeted IP) or quantitative criteria (e.g. fold changes and respective *p* values as determined by statistical testing). For each dataset, we extracted the list of candidate interactors (see Methods for more details) and then examined the overlap between candidate interactors for the same protein from different studies. Surprisingly, the overlap was generally very poor. For example, across four GLH-1 AP-MS experiments, only a single protein meeting the significance threshold replicated across all screens; this protein was GLH-1 itself (Figure 4A). Other proteins show more reproducibility in their identified interactors – ZNFX-1-targeted experiments share four candidates (ZNFX-1, WAGO-4, PRG-1 and LOTR-1; all known Z granule components) (Figure 4B), while MUT-16 experiments share nine (MUT-14, MUT-15, NYN-1, NYN-2, RDE-2, RDE-8, DRH-1, EKL-1, RRF-1, mostly known mutator foci components) (Figure 4C).

**Figure 4.**
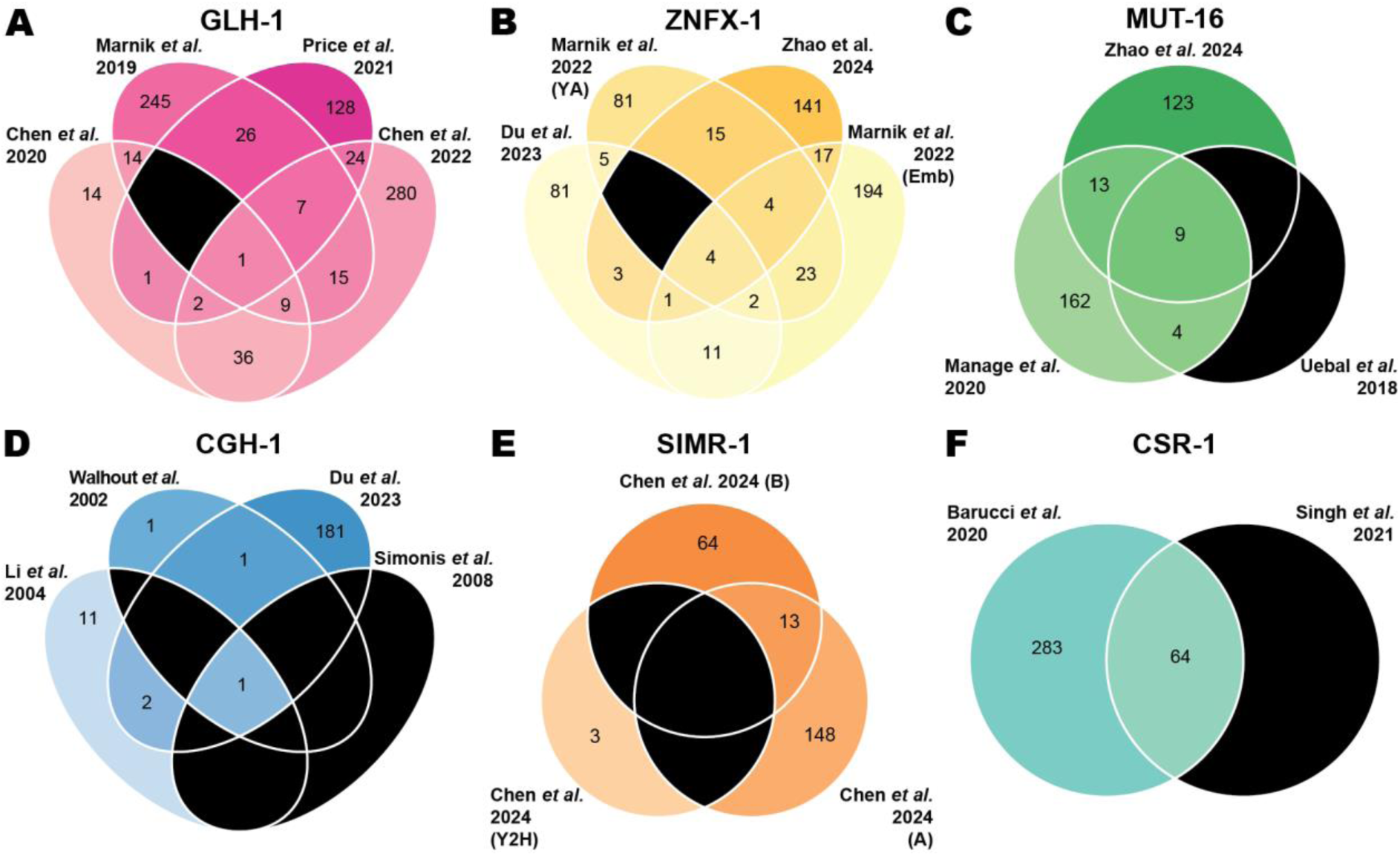
Overlap between significant interactors detected in individual screens. Black segments represent empty intersections. Shown are overlaps between screens for a representative protein from **(A)** P granules (GLH-1), **(B)** Z granules (ZNFX-1), **(C)** Mutator foci (MUT-16), **(D)** P bodies (CGH-1), **(E)** SIMR foci (SIMR-1), and **(F)** D granules (CSR-1).

In some cases, a lack of reproducibility may be explained by differences in the method employed. Three out of four candidate interactor lists for CGH-1 originate from whole-proteome Y2H screens, which may only detect a handful of interactions, while the fourth is derived from an AP-MS experiment; a stark difference in approach and number of identified interactors means that a low level of overlap might be expected (Figure 4D). Likewise, three experiments to identify SIMR-1 interactors, all originating from the same publication (Chen and Phillips 2024), showed no overlapping candidates, however when excluding the Y2H screen and only considering the AP-MS experiments, an overlap is observed (Figure 4E). Some experiments overlap extremely well – one study investigating CSR-1 interactors has all its significant hits shared with a second, independent study (Figure 4F).

### Scoring of proteins across datasets provides more robust indications of their possible granule association

Given the poor overlap between candidates when considering experiments in isolation, we implemented a more holistic approach: an algorithm to search through each dataset and for each detected protein calculate a score for each granule based on the strength of the identification and/or enrichment (see Methods for full details). By adding scores across datasets, the result is a continuous value representing the strength of association between a protein and various granules. Through this method, proteins that are detected robustly across screens are boosted with more sensitivity than a binary significance call as employed for individual experiments.

The resulting matrix (available in Supplemental Table S2) gave each protein a score for each granule examined (7 scores total). When looking at the total sum of all scores for each protein, most proteins had very low total scores, indicating low scores across the board and therefore are likely background proteins with no specific enrichment (Figure 5A). Therefore, we filtered the matrix for only proteins in the top 20% of total scores, with an additional criterion for a minimum of three detections across all experiments analysed. After this filtering step, 920 proteins remained. We performed hierarchical clustering on these proteins to examine how proteins with similar scores grouped together and how this aligned with their potential granule association; to this end we formed 7 clusters, matching the number of granules scored for (Figure 5B). Each cluster contained proteins that mostly had heightened scores for one specific granule, rather than multiple. Cluster 1 contained proteins scoring highly for the E granule, cluster 2 for Mutator foci, cluster 3 for SIMR foci, cluster 4 for the D granule, cluster 5 for P bodies, cluster 6 for P granule, and lastly cluster 7 for Z granule. The full list of proteins in each cluster is available in Supplemental Table S3.

**Figure 5.**
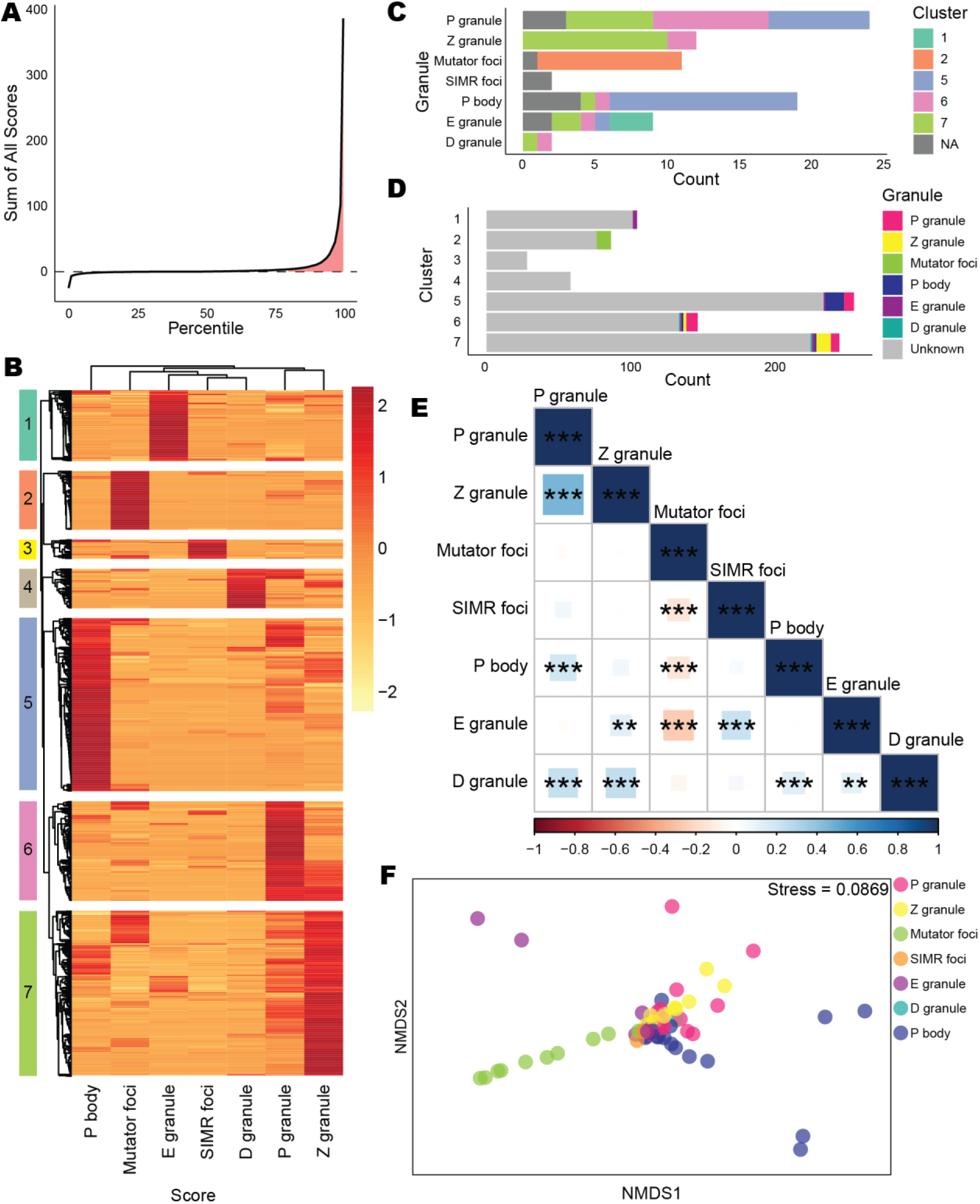
Scoring and clustering of proteins across datasets to group proteins into their potential compartments. **(A)** Percentile distribution of total score sums for all proteins in the score matrix. Shaded in red are the top 20% of proteins, which were taken forward for clustering analysis. **(B)** Clustered heatmap of filtered score matrix. Each row represents a protein, each column represents the score for the indicated granule. Heatmap is scaled by row. Created using the *pheatmap* package using the “average” clustering method (Kolde 2018). **(C)** Bar chart showing where canonical granule components cluster. **(D)** Bar chart showing the distribution of known granule proteins within clusters. **(E)** Spearman correlation matrix of granule scores. Magnitude of correlation coefficient represented graphically using colour and size of square, significant correlations (corrected for multiple comparisons using the Holm method) are indicated on the matrix. * *p* < 0.05, ** *p* < 0.01, *** *p* < 0.001. **(F)** NMDS analysis of score matrix for canonical granule components. Performed using the *vegan* package (Oksanen et al. 2025) with distance set to “manhattan” and dimensions (k) set to 2. Stress < 0.1 indicates good fit (Kruskal 1964).

We then looked at the distribution of proteins between clusters, and how well canonical granule components clustered correctly (Figure 5C). Nineteen canonical P granule components met the detection criteria and were split between three different clusters – clusters 5, 6 and 7. Z granule proteins clustered much more cleanly; of twelve proteins detected, ten were in cluster 7 and two were in cluster 6. All nine canonical Mutator foci components meeting the detection threshold were in cluster 2. Twelve of fifteen detected P body components were in cluster 5, with the remaining three in cluster 6 or 7. Being the most recently discovered granule, the E and D granules had the least amount of data to calculate scores with; nonetheless three of nine canonical E granule components were together in cluster 1, with the remaining six split between cluster 5, 6 or 7. In contrast, neither canonical D granule component clustered together, instead being split between clusters 6 and 7. Neither SIMR foci component met the detection threshold and so were not included in any cluster. Most proteins in each cluster had no known granule-association (Figure 5D).

Relationships between scores for different clusters were further explored with a correlation matrix (Figure 5E). The P granule score showed significant positive correlation with scores for the Z granule, P body and D granule. In contrast, the Mutator foci score showed no positive correlation to any other granule scores, and instead showed a negative correlation with SIMR foci, P body, and E granule scores.

We used non-metric dimensional scaling (NMDS) on the score matrix for the canonical granule components to further examine their variation (Figure 5F). showcases how certain granule components are more distinct than others. P, Z, SIMR and D granule components mostly cluster towards the centre of the NMDS plot, indicating their scores are relatively like each other. Some Mutator foci, E granule and P body components show separation, but this follows a “gradient” of dissimilarity rather than a clean separation.

### Identification of new candidates for granule-associated proteins

As well as providing insights into overall granule architecture, we also believe that the scoring approach could identify candidates for novel granule inhabitants. As such, we searched through the matrix for candidates that showed high association scores for one or more granules but have not had their granule localisation investigated in detail, if at all. Here we provide two examples of proteins that merit further investigation.

The Argonaute PPW-2 was detected in 22 datasets and scored highly for association with the Z granule and P granule, in that order (Figure 6A). PPW-2 has been previously shown to localise to the sperm-specific PEI granule (Schreier et al. 2022), and has also been observed in germ granules in the hermaphrodite germline (Schreier et al. 2022; Seroussi et al. 2023) (Figure 6B). RACK-1 is a scaffolding protein that is known to interact with several proteins and act within several pathways, including signalling, translation regulation, and miRNA function (Nilsson et al. 2004; Jannot et al. 2011). RACK-1 was robustly detected in granule interaction screens (23 detections) and has high association scores for the P granule, Z granule and P body (Figure 6C). RACK-1 shows strong germline cytoplasmic expression, but no clear association with perinuclear germ granules in the hermaphrodite germline (Figure 6D). To date, high-resolution imaging to delineate specific localisation within the germ granules has not been done for PPW-2 nor RACK-1.

**Figure 6.**
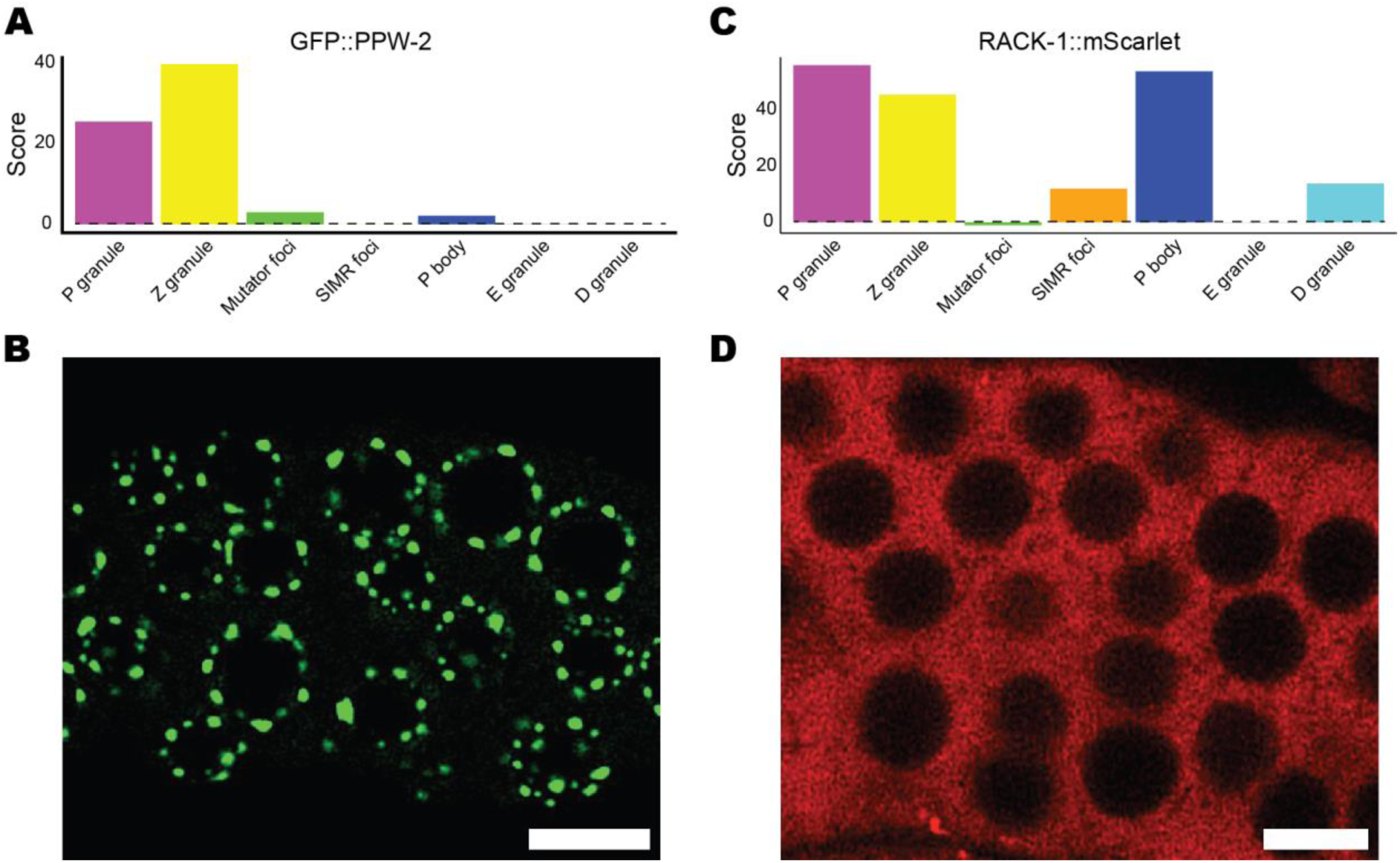
Examples of individual proteins scores and their expression patterns. **(A)** Granule association scores for PPW-2. **(B)** Expression pattern of PPW-2 in the pachytene germline. **(C)** Granule association scores for RACK-1. **(D)** Expression pattern of RACK-1 in the pachytene germline.

## Discussion

### Granule interactome experiments are highly sensitive to technical and biological variation

In this analysis, we found that the reproducibility of significant interactions between individual experiments for the same protein was surprisingly low (Figure 4). This is likely at least partially attributable to experimental and technical variation. In the broadest sense, there are many differences in methods used for PPI detection that would result in expected differences in results. Y2H screens, for example, while allowing for high-throughput analysis of multiple proteins, are missing much of the *in vivo* context that could be critical for protein interaction, such as post-translational modifications and multi-protein networks. However, even when applying a method designed to detect *in vivo* interactions such as AP-MS or proximity labelling, there are many sources of variation. Fluorescent tag choice can change the condensation behaviour of various proteins (Dörner et al. 2026; Fatti et al. 2025); therefore it could be extrapolated that experiments probing the same protein with different tags may be vulnerable to subtle changes in protein condensation affecting native interactions.

Transient or weak interactions may be difficult to detect with AP-MS, depending on the stringency of the washing steps prior to elution of bound proteins. The ability to detect transient interactions is something that proximity labelling techniques seek to address, but these experiments may be prone to higher background and noise, which varies depending on experimental setup and can obscure genuine interactions. Even after protein complexes have been extracted, mass spectrometry acquisition and data analysis introduce further points of potential variation, which contributes to poor reproducibility.

Although technical variation is an undeniable factor, the extreme variance in identified interactions is also reflective of the highly dynamic nature of granules. Some proteins move in, out or between granules depending on certain conditions; for example, HRDE-1 is a nuclear Argonaute (Ashe et al. 2012), but in the absence of its small RNA binding partners can be recruited to SIMR foci by HRDE-2 (Chen and Phillips 2024). Certain P granule and P body components move into stress granules upon cellular stress (Jud et al. 2008). Therefore, experiments that only capture a single point in time may not detect all interactors. Detection of granule-associated proteins may also depend on specific developmental timing. Some Argonaute proteins have been observed to localise to granules, but are only expressed in certain regions of the germline (Seroussi et al. 2023). Likewise, some defined P granule components such as MEG-3 and MEX-3 only localise to granules in the early embryo stage (Wang et al. 2014; Draper et al. 1996). The *in vivo* interaction studies included in this analysis mostly used adult worms, with only a few investigating embryos specifically (Figure 2D). If certain granule components are developmentally or spatially restricted, robustly detecting them could require even further experimental stringency, for example through tight synchronisation of animals or tissue-specific expression of proximity labelling constructs (Holzer et al. 2022).

### Holistic analysis of granule interaction data yields further insight into granule intermixing and interaction

Having observed poor reproducibility between individual experiments, we performed a meta-analysis to take all available data into account. Rather than making binary decisions about whether a protein was significant or not, the scoring algorithm instead provided a continuum indicating the frequency and strength with which each protein associates with a given granule. Hierarchical clustering of the resulting matrix displayed clear groupings aligning with each granule (Figure 5A), suggesting that collective analysis of granule interaction data is at least partially successful in discriminating between different granule compartments. Some granule components appear to cluster very distinctly, most notably the Mutator foci: all its canonical and detected components clustered together, with no known components of other granules. In contrast, some granule components cluster more variably, such as the P granule components, which divided near-equally between clusters 5, 6 and 7 (scoring highly for the P body, P granule or Z granule respectively) (Figure 5B).

There are many explanations for why certain proteins may appear to cluster incorrectly. Some proteins may in fact be misassigned, and in fact localise differently to their currently annotated granule. DEPS-1 and PRG-1 were both originally considered P granule components but were recently found by high resolution microscopy to localise to Z granules (Huang et al. 2024). Therefore, it is not impossible that other proteins have similarly been mischaracterised or show more dynamic localisation than has previously been reported. Notably, in this analysis DEPS-1 and PRG-1 scored highest for Z granules and were in cluster 7 alongside several other Z granule components (Supplemental Table S3), suggesting that our analysis is capable of properly identifying proteins that may be currently misassigned to different granules. Also, while granule sub-compartmentalisation is important for biological function, interaction between granules is possible and oftentimes critical, as has been shown in the case of P granules, Z granules and P bodies (Du et al. 2023). Some Z granule and D granule components have been observed to partially colocalise with P granules, in addition to their primary granule (Huang et al. 2024). Other granule components also display diffuse cytoplasmic expression (Huang et al. 2024) and so may interact with other granules from outside. As such, depending on experimental conditions, protein interaction screens may capture such interactors from outside the specific granule, but are still biologically relevant. Intermixing rather than strict division of granules is also supported by the correlation between scores (Figure 5D). There was significant correlation between the scores for the P granule, Z granule, D granule, and P body – four granules that have been shown to overlap and interact. Furthermore, the NMDS analysis showed that scores for many known granule components are very similar, demonstrated by the points clustering together in the plot (Figure 5E), suggesting their association with distinct granules is not strongly delineated.

In contrast to the high correlation between the aforementioned granule scores, the Mutator foci score does not show strong correlation to other scores (Figure 5D), and many Mutator foci components show distinct separation from other points on the NMDS (Figure 5E). This is somewhat in contrast to previous work that has suggested interaction or partial co-localisation between Mutator foci components and SIMR foci or E granule components (Manage et al. 2020; Chen et al. 2024b; Huang et al. 2024; Phillips et al. 2012). This could suggest a more complex relationship between these granules that has yet to be elucidated. We also noted, however, that the available data probing SIMR foci and E granule interactors is currently very limited (Figure 2C). It is possible that further data on these granules would yield more insight into their relationship with Mutator foci, which may be more in line with previous observations.

Previous efforts to characterise germ granules have provided some insights into their hierarchy. Repeated co-imaging of granules have shown P granules to be at the middle of the germ granule structure, with other granules positioned non-randomly around them (Wan et al. 2018; Du et al. 2023; Zhao et al. 2024; Uebel et al. 2023). Additionally, P granules are the first to be established in the embryo, with subsequent granules appearing as distinct compartments from the 100-cell phase, either arising as a result of demixing from the core P granule (as with the Z granule) (Wan et al. 2018) or from condensing *de novo* at this particular developmental stage (as with the Mutator foci) (Uebel et al. 2020). This information, when considered alongside the high variability in clustering of P granule components supports a model whereby the P granule acts as the “centrepiece” of the germ granule architecture and a hub for facilitating inter-granule interactions. However, P granules are not always required for the formation of other germ granules (Chen et al. 2024b; Phillips et al. 2012), suggesting that they may not serve an essential structural or scaffolding function for broader germ granule architecture.

### Using granule association scores to inform future investigations

While PPI screens are extremely useful for identifying candidate granule residents, microscopy is still the primary method for validating granule association. *C. elegans* is a useful model for this purpose due to the ease of tagging endogenous loci and ability to image live animals at high resolution. Here we have flagged PPW-2 and RACK-1 as two proteins of many that can be mined from the association score analysis (Figure 6). PPW-2 is a known granule inhabitant, but to date its precise localisation in the hermaphrodite germline has not been determined. The granule association scores calculated here suggest that PPW-2 might localise to the Z granule, like other Argonautes such as PRG-1 and WAGO-4 (Wan et al. 2018; Huang et al. 2024). By using the granule association scores to decide which compartments to prioritise checking first, the labour involved in crossing and creating new strains might be minimised.

Scores could also point to candidates that are largely novel in the granule context, such as RACK-1. RACK-1 is a conserved protein with roles in regulating translation (Nilsson et al. 2004) and miRNA function (Jannot et al. 2011). RACK-1 has been shown to interact with the miRNA-induced silencing complex (miRISC), which localises to P bodies (Ding et al. 2005). Furthermore, RACK-1 interacts with the translational repressor GLD-1 (Vanden Broek 2021; Sangari 2022), which shows weak localisation to P granules under standard conditions (Huang et al. 2024; Jones et al. 1996); mutation of *rack-1* increases this localisation dramatically (Vanden Broek 2021). Here, RACK-1 scored highly for association with the P granule, Z granule and P bodies, but preliminary high-resolution imaging did not show distinct perinuclear foci as might be observed in other granule residents. RACK-1 may interact with germ granules from the cytoplasm without concentrating into them itself, perhaps acting as a scaffolder of the germ granule architecture.

Not every high-scoring protein is likely to be a genuine granule inhabitant, but care must be taken when choosing which hits to disregard. For example, ribosomal proteins were some of the most-detected proteins across interaction screens. This is not unusual; ribosomes appear often in mass spectrometry experiments due to their high abundance in the proteome, and may often be disregarded as “junk” hits (Mellacheruvu et al. 2013). In the case of granule interaction studies, however, they may be of interest given the close links between granules, post-transcriptional regulation and translation; ribosomes have even been observed interacting at the interface of granules (Chen et al. 2024a). Therefore, by considering proteins that are consistently detected, we may be able to identify candidates that would otherwise have flown under the radar in an individual experiment.

### Where to next for granule interactome research?

The key limitation to this analysis is the amount of available data. While several interaction screens have been performed for some granules, such as P granule, Z granule, and P body components, other granules are far more limited in their available data (Figure 2C). This is largely explained by time constraints, particularly in the case of the E and D granule that have only been defined in recent years (Chen et al. 2024b; Lu et al. 2025). Experimental complications may also contribute to lack of data – granule proteins may be highly sensitive to changes in structure induced by the addition of a tag (Uebel and Phillips 2019). For example, SIMR-1 function is compromised by the addition of a TurboID epitope, which affected researchers’ ability to probe its interaction partners using this method (Zhao et al. 2024).

Given the relative lack of investigation into SIMR foci, E granules, and D granules, it would be beneficial for future work to prioritise expanding characterisation of these compartments. Through better understanding the proteins that interact within these granules, we can better understand their specific function and how it may intersect with the other, better characterised granules. It would also be prudent to consider how the previously discussed points of technical and biological complication may impact experiments, and ensure that granule interaction investigations are approached from multiple angles: for example, through the use of different tags or methodologies to ensure that identified candidate interactors are robust and reproducible, but also to better capture the dynamic and variable nature of granule proteins. Studies could also look further into how granule interactomes shift in response to development and external stimuli by performing experiments in specific developmental stages and tissues (for example, to better characterise the sperm-specific PEI granule, which was excluded from this analysis due to lack of available data), or under conditions such as those which would lead to the formation of transient stress granules.

Finally, even as the germ granule may continue to be subdivided as more discoveries are made, it is also critical to remember that granules are not isolated, discrete units, but rather a network of intersecting and interacting condensates. Therefore, when detecting interactions for granule proteins, it is important to consider how the results may provide insight not only into the core granule of interest, but also inter-granule associations and interactions. As some proteins have been observed to localise to multiple granules (Chen et al. 2024b; Huang et al. 2024), and even within granules proteins may be spatially arranged in different manners (Wan et al. 2021), future developments in the field may demand a more nuanced and flexible means of referring to and defining what a distinct “granule” really is.

Germ granules and other biomolecular condensates are of rapidly growing interest to research in fields such as cell biology, developmental biology, epigenetics, and more. *C. elegans* will likely continue to maintain its status as a powerful model for exploring their formation, organisation and function, and as such it is important for the field to continue not only generating new data but also consider all current available data to best inform future directions and allow for robust insights to be gained.

## Methods

### Selection and curation of datasets

The literature up to the end of June 2025 was searched for publications that identified interactors of known granule-associated proteins using high-throughput screening methods. We considered papers using yeast two-hybrid (Y2H) screens and/or mass spectrometry (MS) based methods such as AP-MS and TurboID. To find such publications, the WormBase (Sternberg et al. 2024) database was leveraged. Each gene page has an “Interactions” section, which curates identified genetic, physical, and predicted interactions. Interactions were filtered for those annotated as “physical” interactions and accompanying publications were checked for the presence of appropriate experiments. To be eligible for inclusion, publications needed to include the results of the respective experiment: at minimum, a list of proteins identified as candidate interactors in the experiment. In total, 32 publications were found (Table 1).

**Table 1.** Sources of data included in this analysis.

| Publication | Target protein(s) | Target granule(s) |
| --- | --- | --- |
| **Yeast two-hybrid screens** |  |  |
| (Walhout et al. 2002) | Various | Various |
| (Huang et al. 2002) | MEX-3 | P granule |
| (Li et al. 2004) | Various | Various |
| (Simonis et al. 2009) | Various | Various |
| (Chen and Phillips 2024) | SIMR-1 | SIMR foci |
| **Affinity purification-mass spectrometry** |  |  |
| (Audhya et al. 2005) | CAR-1 | P body |
| (Zhang et al. 2007) | AIN-2 | P body |
| (Scheckel et al. 2012) | GLD-1 | P granule |
| (Akay et al. 2013) | GLD-1 | P granule |
| (Zinovyeva et al. 2015) | ALG-1 | P body |
| (Tsai et al. 2015) | RDE-8 | Mutator foci |
| (Chen et al. 2016) | PGL-3 | P granule |
| (Wu et al. 2017) | AIN-1 | P body |
| (Uebel et al. 2018) | MUT-16 | Mutator foci |
| (Marnik et al. 2019) | GLH-1 | P granule |
| (Chen et al. 2020) | GLH-1 | P granule |
| (Manage et al. 2020) | MUT-16 | Mutator foci |
| (Placentino et al. 2021) | PID-2, PID-4, PID-5 | Z granule |
| (Suen et al. 2020) | PRG-1 | Z granule |
| (Barucci et al. 2020) | CSR-1, PRG-1, WAGO-1 | D granule, Z granule, P granule |
| (Cipriani et al. 2021) | MEG-3 | P granule |
| (Singh et al. 2021) | CSR-1, PRG-1 | D granule, Z granule |
| (Marnik et al. 2022) | LOTR-1, ZNFX-1 | Z granule |
| (Cassani and Seydoux 2022) | MEG-1 | P body |
| (Dai et al. 2022) | WAGO-1, PRG-1 | P granule |
| (Chen et al. 2022a) | GLH-1 | P granule |
| (Du et al. 2023) | WAGO-4, ZNFX-1, CGH-1 | Z granule, P body |
| (Chen et al. 2024b) | EGO-1 | E granule |
| **Proximity labelling** |  |  |
| (Price et al. 2021) | GLH-1, DEPS-1 | P granule, Z granule |
| (Zhao et al. 2024) | PGL-1, ZNFX-1, MUT-16 | P granule, Z granule, Mutator foci |
| (Shi et al. 2025) | PRG-1 | Z granule |

### Data curation and quality control

Protein lists and relevant variables were saved from selected publications. The WormBase SimpleMine tool was used to ensure consistency of gene naming. Genes that were not identifiable by SimpleMine were excluded from analysis.

To gather lists of candidate interactors from individual experiments, filtering was applied to the data based on available variables. Some publications merely publish a list of candidate interactors without supplemental data; in this scenario the lists were taken as-is. For large-scale Y2H screens, interactions were often categorised into “core” and “non-core” interactions – only the “core” interactions were considered as significant hits. Mass-spectrometry based experiments typically determine candidate interactors by fold change of protein signal between the target IP and a control IP. Proteins with at least a two-fold change were taken as candidate interactors – if fold changes were precalculated then filtering was based on this, and if not then fold change was calculated from available data. Where *p* values were provided alongside calculated fold change, an additional filter to only include candidates with *p* < 0.05 was applied to ensure appropriate stringency.

### Scoring of proteins to granules

A custom R script was used to calculate granule association scores for each detected protein. In brief, the script looped through each protein in the list of all proteins detected and searched for its presence in each provided dataset. If the protein was detected, +1 was added to a variable keeping count of the number of datasets a protein was detected in. Then, the target of the interaction was retrieved (defined as the bait protein for Y2H experiments, or as the protein being tagged for IP/proximity labelling in the case of AP-MS/TurboID experiments). If the protein being searched for matches the target, the dataset was skipped, as these detections are largely non-informative.

Next, depending on the type of data, a score would be calculated for the protein. For yeast two-hybrid data, the score would be determined based off the confidence of the interaction, and/or the number of times an interaction was detected within the screen. For data where the number of detections was provided, the added score was calculated as a ratio between the number of detections of that specific interaction, and the average number of detections for all interactions detected across the data. Confidence measures were assigned a score from 1-10 (individually tailored to the specific publication’s method of determining confidence, e.g. “core” vs “non-core” or a A-F letter scale). For mass-spectrometry based data, the published data was first searched for already-calculated log_2_(Fold Change), as is commonly reported in such experiments. If this was found, then the score was calculated based on this, with a higher positive fold change compared to the control resulting in a higher score. Where *p* values were provided, interactions were further weighted based on *p* value, with a lower *p* value further enhancing the score. If no *p* values were provided, then no such weighting was performed. If pre-calculated fold changes were unavailable, the algorithm then attempted to use available quantitative data (e.g. LFQ intensity) to calculate a fold change for the protein, which could then be used to add to the score. In the absence of such data, other measures such as number of peptides were utilised to determine a score.

After a score was determined, it was added back to the appropriate granule based on the target of the interaction. For example, if a protein was detected as interacting with PGL-1, PGL-3, and ZNFX-1 with scores for those interactions at 10, 15 and 5 respectively, that protein would receive a final “P granule” score of 25 (10 + 15) and a “Z granule” score of 5. After iterating through every detected protein, the final score matrix was compiled.

### Microscopy

Adults were immobilised on a 2% agarose pad in 5 mM tetramisole, then imaged on a Zeiss LSM800 with Airyscan detector and 63X/1.49 oil objective. Strains used were JMC221 (*ppw-2*(*tor115[GFP::3xFLAG::ppw-2])* I) and JAR50 (*rack-1*(*rns8[rack-1::mScarlet]* IV). GFP was excited with a 480 nm laser and mScarlet was excited with a 561 nm laser. Both lasers were set to 0.4% power and 800 V gain. Images were processed using the inbuilt Airyscan processing at standard power.

### Data processing and visualisation

Unless otherwise stated, all data processing and visualisation was performed using R version 4.4.1 (R Core Team 2024) using tools from the *tidyverse* (Wickham et al. 2019). Euler and Venn diagrams were created using the *eulerr* package (Larsson 2024).

### Code availability

The full R script used to compile the score matrix is available as Supplemental Code.

## Competing interest statement

The authors declare that they have no conflict of interest.

## Acknowledgements

CW was supported by an Australian Government Research Training Program (RTP) Scholarship. AA was supported by DP240100725 from the Australian Research Council. We acknowledge the technical and scientific assistance of Sydney Microscopy & Microanalysis, the University of Sydney node of Microscopy Australia. We also thank the CGC (University of Minnesota), which is funded by NIH Office of Research Infrastructure Programs (P40 OD010440), for providing strains.

## Author contributions

CW and AA conceived and designed the study. CW wrote the code and performed the analysis and microscopy. CW and AA wrote and revised the manuscript. AA supervised and funded the study.

